# Impact of Type II LRRK2 inhibitors on signalling and mitophagy

**DOI:** 10.1101/2021.05.21.445132

**Authors:** Anna Tasegian, Francois Singh, Ian G Ganley, Alastair D Reith, Dario R Alessi

## Abstract

Much effort has been devoted to the development of selective inhibitors of the LRRK2 as a potential treatment for LRRK2 driven Parkinson’s disease. In this study we first compare the properties of Type I (GSK3357679A and MLi-2) and Type II (GZD-824, Rebastinib and Ponatinib) kinase inhibitors that bind to the closed or open conformations of the LRRK2 kinase domain, respectively. We show that Type I and Type II inhibitors suppress phosphorylation of Rab10 and Rab12, key physiological substrates of LRRK2 and also promote mitophagy, a process suppressed by LRRK2. Type II inhibitors also display higher potency towards wild type LRRK2 compared to pathogenic mutants. Unexpectedly, we find that Type II inhibitors, in contrast to Type I compounds, fail to induce dephosphorylation of a set of well-studied LRRK2 biomarker phosphorylation sites at the N-terminal region of LRRK2, including Ser935. These findings emphasize that the biomarker phosphorylation sites on LRRK2 are likely reporting on the open vs closed conformation of LRRK2 kinase and that only inhibitors which stabilize the closed conformation induce dephosphorylation of these biomarker sites. Finally, we demonstrate that the LRRK2[A2016T] mutant which is resistant to MLi-2 Type 1 inhibitor, also induces resistance to GZD-824 and Rebastinib suggesting this mutation could be exploited to distinguish off target effects of Type II inhibitors. Our observations provide a framework of knowledge to aide with the development of more selective Type II LRRK2 inhibitors.

## Introduction

Autosomal dominant mutations that activate the LRRK2 (leucine rich repeat protein kinase-2) signalling pathway cause late onset Parkinson’s disease [1, 2]. LRRK2 is a large 2527 residue multidomain enzyme bearing an armadillo, ankyrin, leucine-rich repeats followed by a tandem Roco type GTPase consisting of a Roc and Cor domain, a Ser/Thr kinase domain and a C-terminal WD40 domain [3]. Various pathogenic mutations have been well-characterized that lie within the LRRK2 Roc (N1437H, R1441G/C/H), Cor (Y1699C), or kinase (G2019S, I2020T) domain [4]. The G2019S mutation is located within the conserved Mg^2+^ subdomain VII motif of the kinase domain and represents the most frequently observed mutation [5]. LRRK2 phosphorylates a subset of Rab GTPases, including Rab8A, Rab10 and Rab12 at a conserved Ser/Thr residue (Thr72 for Rab8A, Thr73 for Rab10, Ser106 for pRab12) [6, 7]. The LRRK2 phosphorylation site is located within the Rab effector-binding Switch-II motif. LRRK2 phosphorylated Rab proteins interact with new set of phospho-binding effectors such as RILPL1 and RILPL2 [6, 8].

All LRRK2 pathogenic mutations increase phosphorylation of Rab substrates including Rab10 [6, 7, 9]. These mutations also stimulate LRRK2 autophosphorylation at Ser1292 [10], but detection of this site is challenging especially for endogenous LRRK2, due to its low stoichiometry of phosphorylation. LRRK2 is also phosphorylated at several well studied serine residues that are termed biomarker sites including Ser910, Ser935, Ser955 and Ser973, located between the ankyrin and leucine rich repeats [11, 12]. Phosphorylation of Ser910 and Ser935 promotes interaction with 14-3-3 scaffolding proteins, preventing LRRK2 from assembling into inclusion like bodies in the cytosol [11, 13]. These biomarker sites have received much attention as they become dephosphorylated following pharmacological inhibition of LRRK2 [13]. Dephosphorylation of Ser935 in particular, has been widely exploited to assess in vivo efficacy of LRRK2 inhibitors [14-16]. It is unclear whether LRRK2 Ser910/Ser935 phosphorylation is regulated by autophosphorylation or via upstream kinases. Phosphorylation of the biomarker sites do not correlate with intrinsic LRRK2 kinase activity as mutation of these residues to Ala do not impact basal LRRK2 kinase activity [11, 17]. Furthermore, various pathogenic mutations including R1441C/G, Y1699C and I2020 mutations suppress the phosphorylation of Ser910 and Ser935 through unknown mechanisms [11, 13].

Evidence points towards LRRK2 controlling mitochondrial [18, 19] and lysosomal [20-24] homeostasis. Recent work has revealed that LRRK2 inhibits basal mitophagy in a variety of cells including dopaminergic neurons and that LRRK2 inhibitors promote mitophagy in LRRK2[G2019S] knock-in mice [25]. LRRK2 has also been shown to control repair of damaged endomembranes through its ability to phosphorylate Rab8A [26, 27]. Treatment of cells with agents that stress lysosomes, promote Rab protein phosphorylation by LRRK2 [28, 29]. Pathogenic LRRK2 also inhibits ciliogenesis in cholinergic interneurons through LRRK2 phosphorylated Rab10 forming a complex with RILPL1 [30, 31]. Cilia loss in cholinergic interneurons induced by LRRK2 pathogenic mutations may decrease ability of these cells to sense Sonic hedgehog in a neuro-protective circuit that supports dopaminergic neurons [30].

There are two widely studied classes of kinase inhibitors namely, Type I that bind to the kinase domain in a closed active conformation and Type II that maintain the kinase in an open inactive-conformation [32]. Most kinase inhibitors are Type I compounds that bind to the ATP binding pocket in a conformation in which the critical catalytic residues including the DFGψ motif (DYGI in LRRK2) are oriented in an “in” conformation with the α-helix and the core hydrophobic spine all aligned in an fully active conformation [33, 34]. Type II inhibitors, include approved kinase compounds such as imatinib [35] and sorafenib [36]. These compounds exploit an additional binding site located adjacent to the region occupied by ATP when the kinase is in the inactive conformation. In this inactive conformation, the DFGψ motif is oriented in an “out “conformation pointing away from the α-helix and in addition the core hydrophobic spine residues are not aligned with each other [32, 37].

R1441G and I2020T pathogenic mutations as well as treatment with Type I LRRK2 kinase inhibitors promote binding of LRRK2 to microtubule filaments [34, 38, 39]. Tomography and cryo-EM analysis suggest that pathogenic mutations induce the LRRK2 kinase domain to adopt a closed active conformation that becomes intrinsically capable of binding microtubules in a well-ordered and periodic manner [40, 41]. Furthermore, recruitment of LRRK2 to microtubules inhibits kinesin and dynein microtubule motility in biochemical studies [41]. High resolution cryoEM studies have recently described the inactive structures of full length [42] as well as the catalytic moiety [43] of LRRK2. Comparison of the active LRRK2 structure predicted from *in situ* Cryo-electron tomography analysis of LRRK2 associated with microtubule filaments [40] as well as AlphaFold artificial intelligence predicted structure of LRRK2 [44], indicates that the active conformation of LRRK2 is markedly distinct from that of inactive LRRK2. Thus, Type I and Type II kinase inhibitors that trap LRRK2 in the inactive vs active conformation may have distinct physiological effects.

Three Type II kinase inhibitors have been reported to target LRRK2 namely GZD-824 [41], Rebastinib [45], and Ponatinib [46], compounds originally developed as inhibitors of the breakpoint cluster region-Abelson (Bcr-Abl) kinase to overcome clinically acquired mutation-induced resistance against imatinib the first line treatment of chronic myelogenous leukemia [43, 47-49]. These compounds also suppressed binding of LRRK2 to microtube filaments, consistent with Type II inhibitors maintaining LRRK2 in an inactive open conformation incapable of interacting with microtubules [41, 45].

In this study we investigate the cellular and invitro properties of Type II LRRK2 inhibitors (GZD-824, Rebastinib and Ponatinib) (Fig 1). We find that Type II inhibitors similar to Type I compounds, suppress LRRK2 mediated phosphorylation of Rab proteins and stimulate basal mitophagy. Unexpectedly, and in contrast to Type I inhibitors, GZD-824, Rebastinib as well as Ponatinib, do not induce the dephosphorylation of LRRK2 at Ser935 and other nearby residues. We also report that Type II inhibitors suppress wild type LRRK2 more potently than pathogenic forms of this enzyme. These observations provide a framework of knowledge that will help with the development and characterization of more selective Type II LRRK2 inhibitors as tools to dissect LRRK2 signaling functions.

**Figure 1.**
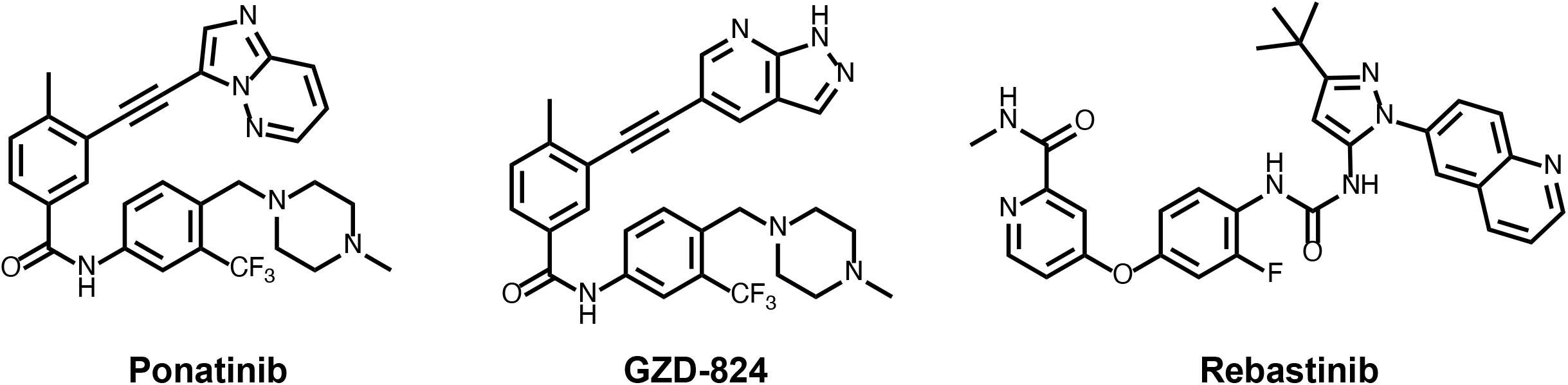
Structures of the LRRK2 Type II inhibitors employed in this study.

## Materials and methods

### Materials

GSK3357679A [25]was synthesized at GlaxoSmithKline. MLi-2 was synthesized by Natalia Shpiro (University of Dundee) as described previously [16]. GZD-824 (#21508) and Rebastinib (#21465) were purchased from Cambridge Bioscience and Ponatinib (#4274) from Tocris. Microcystin-LR was purchased from Enzo Life Sciences (#ALX-350-012-M001). Human recombinant full length wild type Flag-LRRK2 (#A15198) and Flag-LRRK2[G2019S] (#A15201) were from ThermoFisher Scientific. Diisopropylfluorophosphate was from Sigma (#D0879).

### Antibodies

Rabbit monoclonal antibodies for LRRK2 phospho-Ser935 (#UDD2) and phospho-Ser910 (#UDD1) were expressed and purified at University of Dundee as described previously [50]. The C-terminal total LRRK2 mouse monoclonal antibody was from Neuromab (clone N241A/34, #75-253). Rabbit monoclonal antibodies for LRRK2 phospho-Ser1292 (#ab203181), phospho-Ser955 (#ab169521), phospho-Ser973 (#ab181364), and the recombinant MJFF-phospho-Thr73-Rab10 (#ab230261) [51], MJFF-Rab10 total (#ab237703), MJFF-phospho-Ser106-Rab12 (#ab256487) and MJFF-phosphoThr71-Rab29 (#ab241062) rabbit monoclonal antibodies were purchased from Abcam. The total Rab10 mouse antibody was from nanoTools (0680–100/Rab10-605B11, www.nanotools.de) [51]. The total Rab12 sheep polyclonal antibody was generated at University of Dundee, MRC PPU Reagents and Services (1^st^ bleed, #SA224). Total Rab12 rabbit polyclonal antibody was from Proteintech (#18843-1-AP). Anti HA rat monoclonal antibody was purchased from Sigma-Roche (clone 3F10, #11867423001). The anti (pan)14-3-3 rabbit polyclonal antibody (#8312) and anti α-tubulin mouse monoclonal antibody (clone DM1A, mAb #3873) were purchased from Cell Signaling Technology. Anti-glyceraldehyde-3-phosphate dehydrogenase (GAPDH) antibody was from Santa Cruz Biotechnology (#sc-32233).

### Plasmids and transient transfections

cDNA constructs for overexpression of human LRRK2 and Rabs employed in the present study were the following: FLAG-LRRK2-WT (DU6841), HA-Rab29 (DU50222), HA-empty (DU44059), GFP-LRRK2 WT (DU13633), GFP-empty (DU13156) and were generated at the University of Dundee, MRC PPU Reagents and Services and are available on request (https://mrcppureagents.dundee.ac.uk/). Plasmids were amplified in *Escherichia coli* DH5a and purified using HiSpeed Plasmid Maxi Kit (Qiagen).

### Cell cultures, treatments, and lysis

Wild-type and homozygous LRRK2 [R1441C] (Jax, stock number: 009346) [6], LRRK2[G2019S] Taconic, model 13940) [7] and LRRK2 [A2016T] (Jax, stock number: 021828)[7] knock-in MEF cells were isolated from littermate-matched mouse embryos on a C57BL/6J background at day E12.5 as described previously. MEFs were grown in Dulbecco’s modified Eagle medium (DMEM GIBCO, #11960-085) containing 10% (by vol) fetal bovine serum (FBS), 2 mM L-glutamine, 100 U/ml penicillin, and 100 mg/ml streptomycin supplemented with non-essential amino acids and 1 mM sodium pyruvate. Human embryonic kidney 293 (HEK293) (#ATCC CRL-1573) and HeLa (#ATCC CCL-2) cells were purchased from American Tissue Culture Collection (ATCC) and cultured in DMEM containing 10% (by vol) FBS, 2 mM L-glutamine, 100 U/ml penicillin, and 100 mg/ml streptomycin. All cells were grown at 37°C, 5% (by vol) CO_2_ in a humidified atmosphere and regularly tested for mycoplasma contamination. The wild type and homozygous LRRK2[G201S] human lymphoblastoid cell lines were described previously [13] and maintained in suspension culture in Roswell Park Memorial Institute (RPMI) 1640 medium with 10% FBS, 2 mM glutamine, 100 U/ml penicillin, and 100 mg/ml streptomycin at cell density of 0.3 × 10^6^-2 × 10^6^ cells per ml. For experiments cultures of 3.5-5 million cells were utilized.

### Inhibitor treatments

Inhibitors were dissolved in dimethyl sulfoxide (DMSO) (Sigma, #D2650). The inhibitors were diluted 1000x into cell medium. The equivalent volume of DMSO was added to the minus inhibitor controls. Experiments were performed in 6-well plates, 10 cm dishes,15 cm dishes or 15 ml Falcon-like tubes depending on the amount of cell extract required.

### Transient transfections

Transient transfections were performed for 24 h using the linear polyethylenimine (PEI, Polyscience, #24765) method [52] with a final concentration of 2 µg/ml PEI and 6 μg of DNA for a 10 cm diameter dish with 10 ml of medium.

### Cell Lysis

Cells were lysed in an ice-cold lysis buffer containing 50 mM Tris–HCl, pH 7.4, 1% (by vol) Triton X-100, 10% (by vol) glycerol, 150 mM NaCl, 1 mM sodium orthovanadate, 50 mM NaF, 10 mM 2-glycerophosphate, 5 mM sodium pyrophosphate, 1 µg/ml microcystin-LR, and cOmplete EDTA-free protease inhibitor cocktail (Roche, #11836170001). Lysates were incubated on ice for 30 minutes and clarified by centrifugation at 20800 x g at 4°C for 20 min. Supernatants were quantified by Bradford assay (Bio-Rad Protein Assay Dye Reagent Concentrate #5000006) and samples prepared in 1x lithium dodecyl sulfate sample buffer final concentration (NuPage LDS Sample buffer 4x Invitrogen, #NP0008). When not deployed immediately, lysates were snap-frozen in liquid nitrogen and stored at ™80 °C until use.

### IC_50_ LRRK2 Nictide assay

Reactions were undertaken with wild type Flag-LRRK2 (#A15198) and Flag-LRRK2[G2019S] (#A15201) purchased from ThermoFisher Scientific. Peptide kinase assays, performed in triplicate, were set up in a total volume of 25.5 μL containing 13.7 ng LRRK2 kinase in 50 mM Tris/HCl, pH 7.5, 0.1 mM EGTA, 10 mM MgCl_2_, 20 μM Nictide (an optimized peptide substrate for LRRK2 [53]), 0.1 μM [γ-^33^P]ATP (∼500 cpm/pmol) and the indicated concentrations of inhibitor dissolved in DMSO. After incubation for 30 min at room temperature °C, reactions were terminated by harvesting the whole reaction volume onto P81 phosphocellulose filter plate and immersion in 50 mM phosphoric acid. Samples were washed extensively and the incorporation of [γ-^33^P]ATP into Nictide was quantified by Cerenkov counting using a Tri-Carb 4910 TR PerkinElmer liquid scintillation analyzer. IC_50_ values were calculated with GraphPad Prism using non-linear regression analysis.

### Kinase Profiling

Protein kinase profiling of the Type I (GSK3357679A and MLi-2) and Type II inhibitors (GZD-824 and Ponatinib) were undertaken at a concentration of 0.1 and 1 μM and carried out against the Dundee panel of 140 protein kinases at the International Centre for Protein Kinase Profiling (http://www.kinase-screen.mrc.ac.uk). The kinase profiling of each inhibitor was undertaken at a concentration of 1 and/or 10 μM. Results for each kinase are presented as the mean kinase activity ± S.D. for an assay undertaken in duplicate relative to a control kinase assay in which the inhibitor was omitted. Abbreviations and assay conditions used for each kinase are available (http://www.kinase-screen.mrc.ac.uk/services/premier-screen).

### Quantitative immunoblot analysis

Cell extracts were mixed with a quarter of a volume of NuPage LDS Sample buffer 4x Invitrogen and heated at 95 °C for 5 minutes. 10-30 μg of samples was loaded onto NuPAGE 4–12% Bis–Tris Midi Gel (Thermo Fisher Scientific, Cat# WG1403BOX) and electrophoresed with the NuPAGE MOPS SDS running buffer (Thermo Fisher Scientific, Cat# NP0001-02) at 90 V. After 30 min the voltage was increased at 120 V for 1 h 30 min. At the end of electrophoresis, proteins were electrophoretically transferred onto the nitrocellulose membrane (GE Healthcare, Amersham Protran Supported 0.45 µm NC) at 100 V for 90 min on ice in the transfer buffer (48 mM Tris–HCl and 39 mM glycine supplemented with 20% methanol). Transferred membrane was blocked with 5% (w/v) skim milk powder dissolved in TBS-T [20 mM Tris–HCl, pH 7.5, 150 mM NaCl and 0.1% (v/v) Tween 20] at room temperature for 1 h. The membrane was typically cropped into three pieces, namely the ‘top piece’ (from the top of the membrane to 75 kDa), the ‘middle piece’ (between 75 and 30 kDa) and the ‘bottom piece’ (from 30 kDa to the bottom of the membrane). All primary antibodies were used at 1 μg/ml final concentration and incubated in TBS-T containing 5% (by mass) bovine serum albumin with exception of α-tubulin and GAPDH antibodies that were diluted 1:5000 and 1:2000 respectively. The top piece was incubated with the indicated rabbit anti-LRRK2 phosphorylation site antibody multiplexed with mouse anti-LRRK2 C-terminus total antibody. The middle piece was incubated with mouse anti-GAPDH or anti-a-tubulin antibody. The bottom pieces were incubated with indicated total and phospho-Rab antibodies. Membranes were incubated in primary antibody overnight at 4°C. Prior to secondary antibody incubation, membranes were washed three times with TBS-T for 10 min each. The top and bottom pieces were incubated with goat anti-mouse IRDye 680LT (#926-68020) secondary antibody multiplexed with goat anti-rabbit IRDye 800CW (#926-32211) secondary antibody diluted in TBS-T (1:25000 dilution) for 1 h at room temperature. The middle piece was incubated with goat anti-mouse IRDye 800CW (#926-32210) secondary antibody diluted in TBS-T (1:25000 dilution) at room temperature for 1 h. Membranes were washed with TBS-T for three times with a 10 min incubation for each wash. Protein bands were acquired via near infrared fluorescent detection using the Odyssey CLx imaging system and quantified using the Image Studio software.

### Analysis of 14-3-3 binding

This was undertaken as described previously [11]. Briefly, wild type FLAG-LRRK2 was transiently expressed in HEK293 cells with HA-Rab29. 24h post-transfection cells were lysed with lysis buffer and snap frozen. LRRK2 was immunoprecipitated from 4 mg of lysate using 20 μl anti-FLAG-M2 agarose beads (Sigma, #A2220) for 2 h in rotation at +4°C. Immunoprecipitates were washed once with lysis buffer supplemented with 0.5% (by vol) Nonidet P-40 (instead of Triton X-100) and 0.5 M NaCl followed by two washes. Samples were eluted in 2x LDS sample buffer (20 µl per 10 µl of resin) and heated at 70 °C for 10 min. After centrifugation through 0.22 µm Spin X filter, the eluted protein was subjected to immunoblotting with the indicated antibodies. 10 μl of the final reaction mixture was needed for the detection of binding of endogenous 14-3-3 and 1 µl for detection of LRRK2.

### Immunofluorescence microscopy

HeLa cells were seeded on coverslips and transfected with the plasmid reported in each image 24 h before fixation. Following the treatments with the inhibitors described in figure legends, cells were washed twice in DPBS and fixed with 3.7% Paraformaldehyde pH=7.3 (Agar scientific, #R1026), 200 mM HEPES, for 20 minutes. Cells were then washed twice with DMEM, 20mM HEPES, and then incubated for 10 minutes with DMEM, 10mM HEPES. After two washes in DPBS, fixed cells were permeabilized with 0.2% (by vol) NP-40 in DPBS for 10 min. Cells were blocked using 1% (by mass) BSA in DPBS, then incubated for 1 h with primary antibodies diluted in 1% (by mass) BSA in DPBS, washed three times in 0.2% (by mass) BSA in DPBS, and incubated for 1 h with secondary antibodies. The coverslips were washed three times with 0.2% (by mass) BSA in PBS. Coverslips were washed once more in double distilled water and mounted on slides (VWR, #631-0909) using ProLong™ Diamond Antifade Mountant (ThermoFisher, #P36961). For localization of the Golgi compartment, Golgin-97 was stained with rabbit anti-GM97 (CST, #D8P2K) and visualized with goat anti-rabbit conjugated to Alexa Fluor 405 (Life Technologies, #P10994). HA-Rab29 was stained with mouse anti-HA (Abcam, clone HA.C5 #ab18181) and visualized with goat anti-mouse conjugated to Alexa Fluor 594 (Life Technologies, #A-11032). The images were collected on an LSM710 laser scanning confocal microscope (Carl Zeiss) using the Å∼63 Plan-Apochromat objective (NA 1.4).

### Human neutrophils isolation

Neutrophils were isolated from 20 ml of whole blood collected from two healthy donors using an immune-magnetic negative isolation described previously [54]. 5 million cells from each condition were re-suspended in 1 ml of RPMI medium at and treated the indicated inhibitors or DMSO as a control for 45 min at room temperature. Cells were pelleted by centrifugation at 335xg for 5 min, the supernatants decanted, and the pellets lysed in lysis buffer freshly supplemented with 0.5 mM diisopropylfluorophosphate to inhibit high levels of intrinsic serine protease activity present in neutrophils [54]. Neutrophil lysates were incubated on ice for 30 minutes, centrifuged at 20800 x g for 20 min at 4 °C, snap-frozen in liquid nitrogen and stored at -80 °C. The human biological samples were sourced ethically, and their research use was in accord with the terms of the informed consents under an IRB/EC approved protocol.

### Mito-QC primary mouse embryonic fibroblasts culture

To assess mitophagy, wild type, G2019S knock in, and LRRK2 knock out primary mouse embryonic fibroblasts (MEFs) were derived, from time-mated pregnant females at E12.5 as previously described [25]. Briefly, cells were plated on glass coverslips and incubated with DMEM (Gibco, 11960-044) supplemented with 10% FBS, 2 mM L-Glutamine (Gibco, 2503-081), 1% non-essential amino acids (Gibco, 11140-035), 1% Antibiotics (Penicillin/Streptomycin 100 U/ml penicillin and 100 μg/ml streptomycin; Gibco), and 150μM β-Mercaptoethanol (Gibco, 21985-023) at 37ºC under a humidified 5% CO2 atmosphere. MEFs were treated for 24 hours with GZD-824 (3, 10, 30, 100, 300 nM). GSK3357679A 100nM (Ding et al., in prep) was used as a positive control. At the end of the treatment, MEFs were briefly washed twice in DPBS (Gibco, 14190-094), and fixed in 3.7% Paraformaldehyde pH=7.00 (Sigma, P6148), 200 mM HEPES, for 20 minutes. Cells were then washed two times with DMEM, 10mM HEPES, and then incubated for 10 minutes with DMEM, 10mM HEPES. Cells were washed in DPBS and mounted on a slide (VWR, Superfrost, #631-0909) with Prolong Diamond (Thermo Fisher Scientific, #P36961). Images were acquired using a Nikon Eclipse Ti-S fluorescence microscope with a 63x objective. Images were analyzed using the mito-QC counter as previously described [55]. Experiments were performed in 3-4 independent experiments and data was analyzed with a one-way ANOVA followed by a Dunnett’s multiple comparison test using the GraphPad Prism software (version 8.4.3 (686)).

## Results

### Type-II inhibitors target wild type LRRK2 with greater potency than pathogenic LRRK2[G2019S] in vitro

We first assessed the in vitro potency of GZD-824, Rebastinib and Ponatinib towards full length wild type and G2019S pathogenic LRRK2 mutant in a biochemical assay using the synthetic peptide substrate termed Nictide [53] at an ATP concentration of 100 μM. This revealed that GZD-824 inhibited wild type LRRK2 and LRRK2[G2019S] with an IC_50_ of 17 nM and 80 nM respectively. Rebastinib inhibited wild type LRRK2 and LRRK2[G2019S] with and IC_50_ of 192 nM and 737 nM respectively. Ponatinib targeted wild type LRRK2 (IC_50_ of 100 nM) ∼4-fold more potently than LRRK2[G2019S] (IC_50_ of 400 nM) (Table 1). The two Type I LRRK2 kinase inhibitors used in this study, namely GSK3357679A and MLi-2, inhibited LRRK2 with sub-nanomolar potency with MLi-2 inhibiting LRRK2[G2019S] ∼2-fold more strongly than wild type LRRK2. GSK3357679A displayed similar potency towards wild type LRRK2 and LRRK2[G2019S] (Table 1).

**Table 1.**
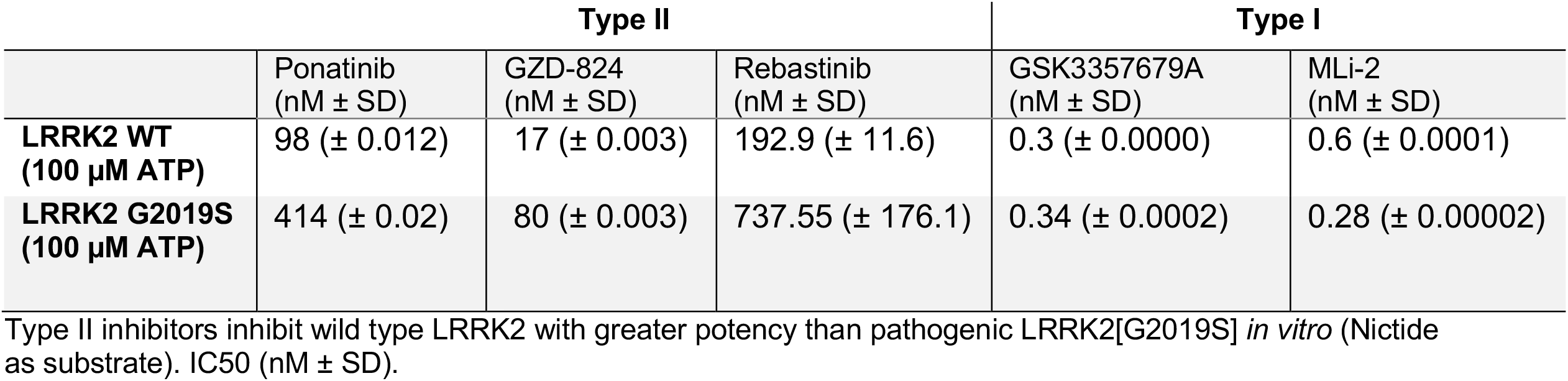
Type-II inhibitors inhibit wild type LRRK2 with greater potency than pathogenic LRRK2[G2019S] in vitro. IC_50_ values the indicated inhibitors towards wild type and LRRK2[G2019S]. LRRK2 assays were undertaken deploying the Nictide peptide substrate [53].

|  | Type II |  |  | Type I |  |
| --- | --- | --- | --- | --- | --- |
| | Ponatinib<br>(nM ± SD) | GZD-824<br>(nM ± SD) | Rebastinib<br>(nM ± SD) | GSK3357679A<br>(nM ± SD) | MLi-2<br>(nM ± SD) |
| **LRRK2 WT**<br>**(100 μM ATP)** | 98 (± 0.012) | 17 (± 0.003) | 192.9 (± 11.6) | 0.3 (± 0.0000) | 0.6 (± 0.0001) |
| **LRRK2 G2019S**<br>**(100 μM ATP)** | 414 (± 0.02) | 80 (± 0.003) | 737.55 (± 176.1) | 0.34 (± 0.0002) | 0.28 (± 0.00002) |

Type II inhibitors inhibit wild type LRRK2 with greater potency than pathogenic LRRK2[G2019S] *in vitro* (Nictide as substrate). IC50 (nM ± SD).

We also assessed the selectivity of GZD-824 (Fig S1A), Rebastinib (Fig S1B) and Ponatinib (Fig S1C) towards a panel of 140 protein kinases. This revealed that all Type II inhibitors were not selective for LRRK2 and inhibited other serine/threonine as well as tyrosine kinases. At 0.1 and 1 μM, GZD-824 suppressed the activity of 22 and 43 of the 140 kinases assayed by over 80% respectively (Fig S1A). 1 μM Rebastinib suppressed 18 out of 140 kinases in the panel by more than 80%. 0.1 and 1 μM Ponatinib suppressed 14 and 20 kinases respectively by 80%. By contrast, GSK3357679A (Fig S1D) and MLi-2 [16] are highly selective kinase inhibitors.

### Type-II LRRK2 inhibitors suppress LRRK2 mediated Rab10 phosphorylation without suppressing Ser935 phosphorylation

We next assessed the ability of GZD-824 (Fig 2A), and Ponatinib (Fig 2B) to inhibit LRRK2 mediated phosphorylation of Rab10 at Thr73 in LRRK2 wild type mouse embryonic fibroblast cells (MEFs). Cells were pre-incubated for 2 h with the indicated doses of GZD-824 or DMSO as a control and phosphorylation of Rab10 (Thr73), Rab12 (Ser105) and LRRK2 at Ser935 monitored by quantitative phospho-immunoblotting. GZD-824 induced a dose dependent inhibition of Rab10 and Rab12 phosphorylation with an IC_50_ of ∼300 nM (Fig 2A). 1 μM GZD-824 reduced Rab10 and Rab12 phosphorylation to almost background levels like that observed with the MLi-2 Type I Inhibitor [16]. Strikingly, GZD-824 did not induce dephosphorylation of LRRK2 at Ser935, under conditions which MLi-2 induced complete dephosphorylation of this site (Fig 2A). Increasing GZD-824 concentrations to 10 μM did not reduce LRRK2 phosphorylation at Ser935 significantly (SFig2A). We observed that concentrations of 10 μM or higher GZD-824 impacted cell morphology and attachment, so we employed GZD-824 at 5 µM or less in subsequent studies. Ponatinib was a weaker inhibitor, and only partially reduced Rab10 phosphorylation ∼50% inhibition at 10 μM, also without affecting Ser935 phosphorylation (Fig 2B). Concentrations of Ponatinib above 10 μM were also observed to impact cell morphology and attachment. GZD-824 suppressed Rab10 and Rab12 phosphorylation in a time dependent manner, with a 50% reduction observed within ∼10 min (Fig 2C). Rab10 and Rab12 phosphorylation remained suppressed and Ser935 phosphorylation unaffected over a 24 h time course of 1 μM GZD-824 treatment (Fig 2D). GZD-824 also inhibited Rab10 phosphorylation in a dose dependent manner in primary human neutrophils isolated from two donors with maximal inhibition observed at 300 nM GZD-824 (Fig 3). Phosphorylation of Ser935 as well as phosphorylation of other biomarker sites Ser910, Ser955 and Ser973 was unaffected by concentration of GZD-824 up to 3 μM (Fig 3).

**Figure 2.**
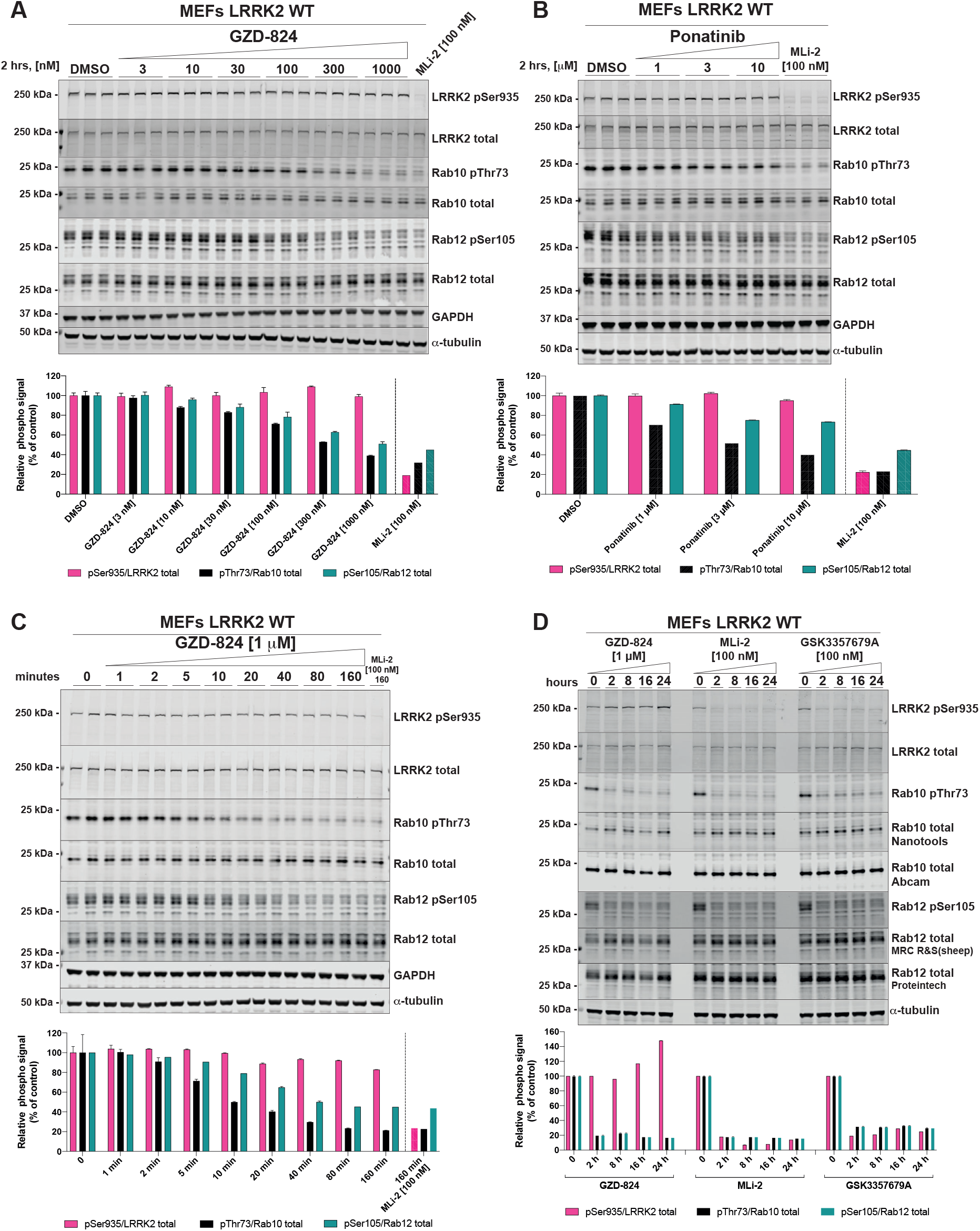
GZD-824 suppresses LRRK2 mediated Rab10 phosphorylation without inhibiting Ser935 phosphorylation. (A+B) Wild-type MEFs were treated with or without the indicated concentrations of inhibitors for 2 h. Cells were lysed, and 20 µg of extract was subjected to quantitative immunoblot analysis with the indicated antibodies (all at 1 µg/ml). Each lane represents cell extract obtained from a different dish of cells (three replicates per condition). The membranes were developed using the Odyssey CLx Western Blot imaging system. (C) As in (A) except that cells were treated with 1 μM GZD-824 or 100 nM MLi-2 for the indicated times. (D) As in (A) except that cells were treated with 1 μM GZD-824 or 100 nM MLi-2 or 100 nM GSK3357679A for 24 h. (A to D) Immunoblots were quantified using the Image Studio software. Data are presented relative to the phosphorylation ratio observed in cells treated with DMSO (no inhibitor), as mean ± SD. (A to C) SD are derived from the replicates show in in the presented blots. (D) SD are derived from duplicates run in independent gels.

**Figure 3.**
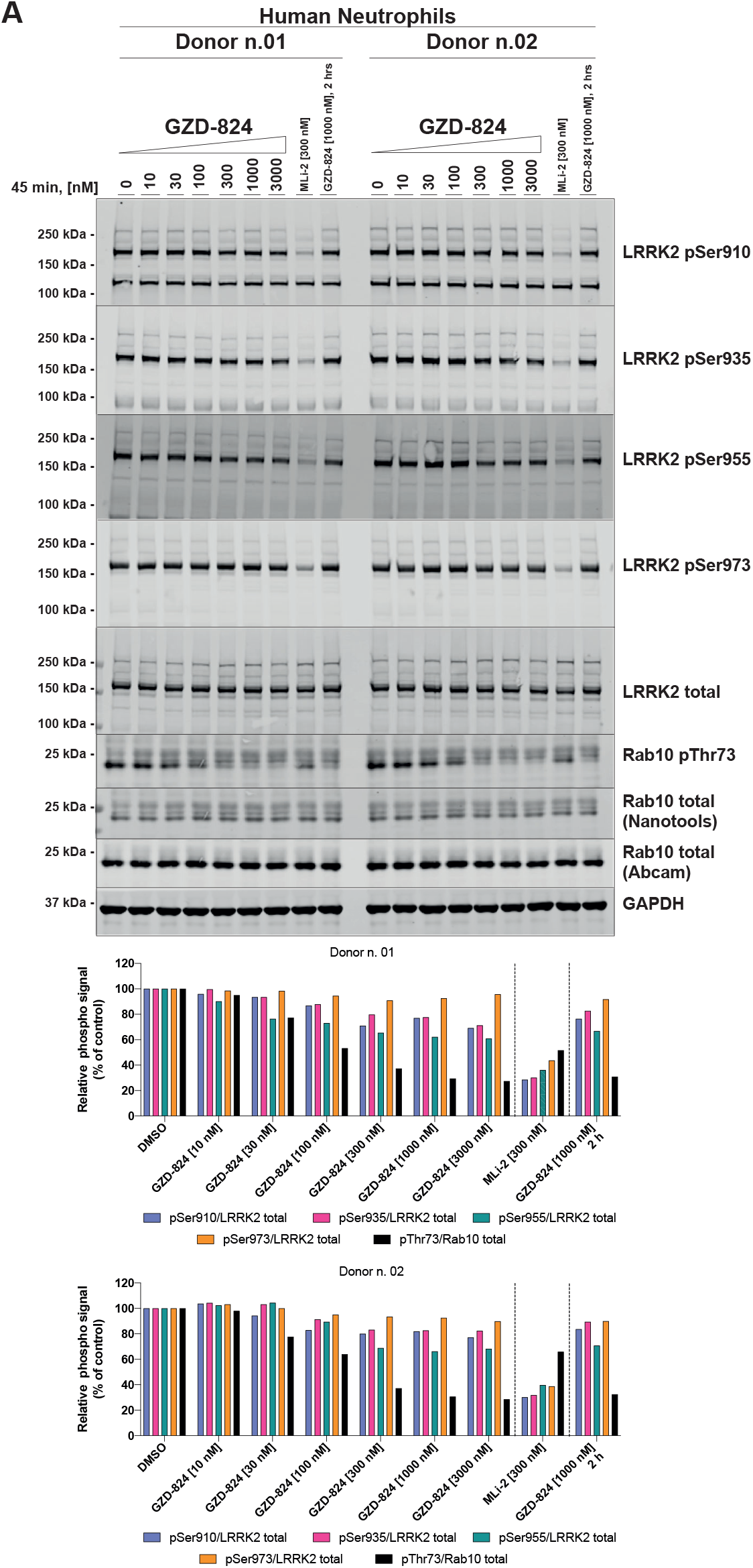
GZD-824 does not inhibit Ser935 phosphorylation in neutrophils. (A) Primary neutrophils from two healthy donors were isolated as described previously [54], and treated with or without 1 μM GZD-824 or 300 nM MLi-2 for 45 min at ambient temperature for the indicated times. Cells were lysed and 25 µg of extract was subjected to quantitative immunoblot analysis with the indicated antibodies (all at 1 µg/ml). The membranes were developed using the Odyssey CLx Western Blot imaging system. Immunoblots were quantified for phospho-Thr73 Rab10/total Rab10 ratio, phospho-Ser935 LRRK2/total LRRK2 ratio using the Image Studio software. Data are presented relative to the phosphorylation ratio observed in cells treated with DMSO (no inhibitor).

### GZD-824 and Rebastinib inhibit pathogenic LRRK2[R1441C] and LRRK2[G2019S] with lower potency than wild type LRRK2

We next compared the potency at which GZD-824 suppressed Rab10 phosphorylation in wild type and LRRK2[R1441C] (Fig 4A) as well as LRRK2[G2019S] knock-in MEFs (Fig 4B). In these experiments we observed that higher doses of GZD-824 were required to reduce phosphorylation of Rab10 in the pathogenic mutant cells. We estimate that the IC_50_ for dephosphorylation of Rab10 in wild type, R1441C and G2019S MEFs is 70 nM, 262 nM, and 233 nM respectively. 1 μM GZD-824 reduced Rab10 phosphorylation to almost basal levels in wild type cells, but only reduced Rab10 phosphorylation ∼ 3 to 3.5-fold in LRRK2[R1441C] MEFs and LRRK2[G2019S] MEFs. Earlier work revealed that the LRRK2[R1441C] mutation reduces LRRK2 Ser935 phosphorylation [11]. Consistent with this, we observed reduced phosphorylation of LRRK2 at Ser935 in the LRRK2[R1441C] knock-in cell lines (Fig 4A, 4C, 4D). Treatment with GZD-824 rescued Ser935 phosphorylation in LRRK2[R1441C] knock-in cells in a dose (Fig 4A) and time dependent manner (Fig 4C, 4D) to levels similar to that observed in wild type cells (Fig 4A, 4C, 4D). Time course analysis revealed that 1 μM GZD-824 increased Ser935 phosphorylation in LRRK2[R1441C] cells by 40 min plateauing at 160 min (Fig 4C, 4D). 1 µM GZD-824 only partially reduced by ∼2-fold Rab10 phosphorylation in LRRK2[R1441C] knock-in cells (Fig 4C, 4D). As reported previously [11], the G2019S mutation did not reduce Ser935 phosphorylation and treatment of these cells with GZD-824 did not impact Ser935 phosphorylation (Fig 4B).

**Figure 4.**
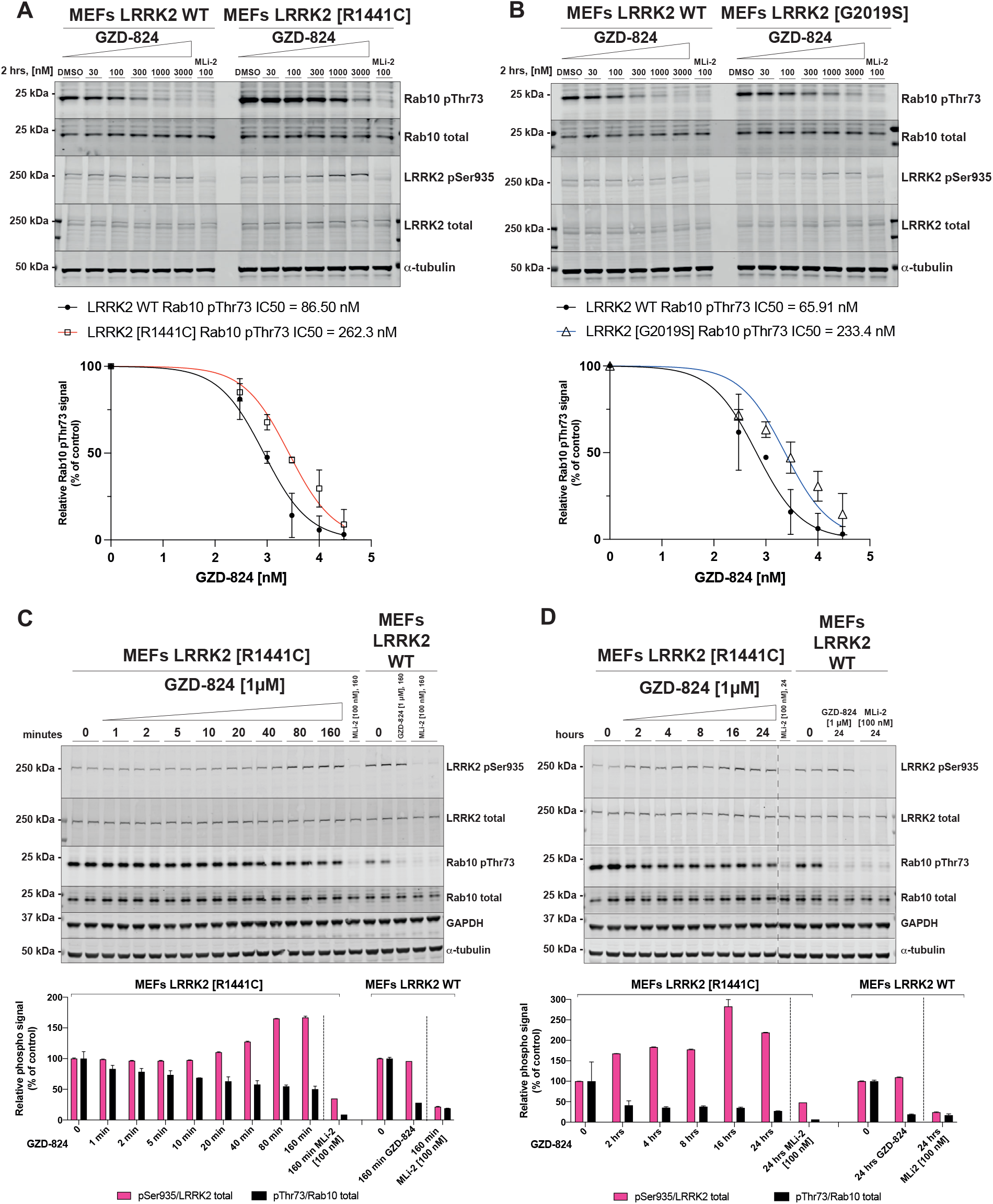
GZD-824 inhibits pathogenic LRRK2 with lower potency than wild type LRRK2. (A+B) The indicated littermate matched wild type and LRRK2 pathogenic knock-in MEFs (passages n. 15-16) were treated with or without the indicated concentrations of inhibitors for 2 h. Cells were lysed, and 25 µg of extract was subjected to quantitative immunoblot analysis with the indicated antibodies (all at 1 µg/ml). Each lane represents cell extract obtained from a different dish of cells. The membranes were developed using the Odyssey CLx Western Blot imaging system. (C+D) As in (A) except that cells were treated with 1 μM GZD-824 or 100 nM MLi-2 for the indicated times. Each lane represents cell extract obtained from a different dish of cells (two replicates per condition). (A to D) Immunoblots were quantified using the Image Studio software. Data are presented relative to the phosphorylation ratio observed in cells treated with DMSO (no inhibitor), as mean ± SD. (A+B) SD are derived from duplicates run in independent gels. (C+D) SD are derived from the replicates show in the presented blots. IC_50_ values were calculated with GraphPad Prism (version 9.1.0) using non-linear regression analysis.

We also compared the ability of Rebastinib to suppress the activity of Rab10 phosphorylation in wild type and LRRK[G2019S] (Fig 5A) and LRRK2[R1441C] MEFs (Fig 5B). Like GZD-824, Rebastinib inhibited wild type LRRK2 with ∼2-fold higher potency than pathogenic mutations. We also observed that Rebastinib like other Type II inhibitors tested did not induce dephosphorylation of Ser935 despite efficiently inhibiting Rab10 phosphorylation (Fig 5A, 5B).

**Figure 5.**
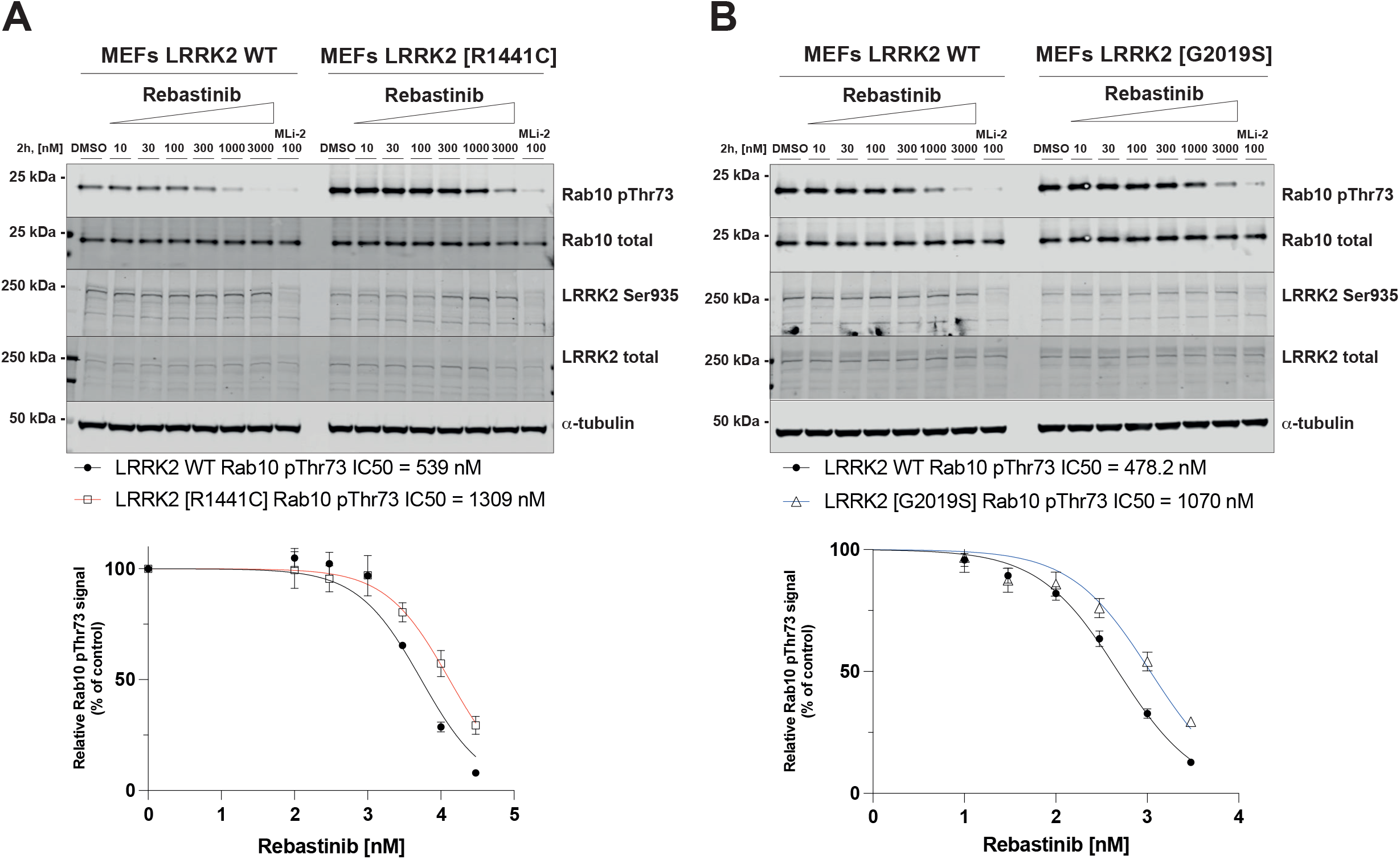
Rebastinib inhibits pathogenic LRRK2 with lower potency than wild type LRRK2. (A+B) The indicated littermate matched wild type and LRRK2 pathogenic knock-in MEFs (passages n. 15-16) were treated with or without the indicated concentrations of inhibitors for 2 h. Cells were lysed, and 25 µg of extract was subjected to quantitative immunoblot analysis with the indicated antibodies (all at 1 µg/ml). Each lane represents cell extract obtained from a different dish of cells. The membranes were developed using the Odyssey CLx Western Blot imaging system. Immunoblots were quantified using the Image Studio software. Data are presented relative to the phosphorylation ratio observed in cells treated with DMSO (no inhibitor), as mean ± SD. (A+B) SD are derived from duplicates run in independent gels. IC_50_ values were calculated with GraphPad Prism (version 9.1.0) using non-linear regression analysis.

### The MLi-2 inhibitor resistant LRRK2 [A2016T] mutation suppresses sensitivity to GZD-824 and Rebastinib

In previous work we have characterized a mutation within the kinase domain of LRRK2[A2016T], that does not impact on catalytic activity but reduced around 10-fold sensitivity towards certain Type I inhibitors tested including MLi-2 [7, 11, 13]. This mutation has been exploited to confirm physiological responses induced by MLi-2 such as Rab protein phosphorylation are indeed mediated by LRRK2 rather than an off-target effect of the inhibitor [7]. We therefore tested whether the A2016T mutation would impact sensitivity towards GZD-824 (Fig 6A, 6B) and Rebastinib (Fig 6C, 6D) using previously described wild type and LRRK2[A2016T] knock-in MEFs [7]. These studies revealed that the A2016T mutation markedly blocked the ability of both GZD-824 and Rebastinib to inhibit Rab10 phosphorylation, even at 3 µM, the highest concentration studied (Fig 6). As expected, the LRRK2[A2016T] mutation also markedly suppressed inhibition of Rab10 phosphorylation induced by the Type I MLi-2 inhibitor.

**Figure 6.**
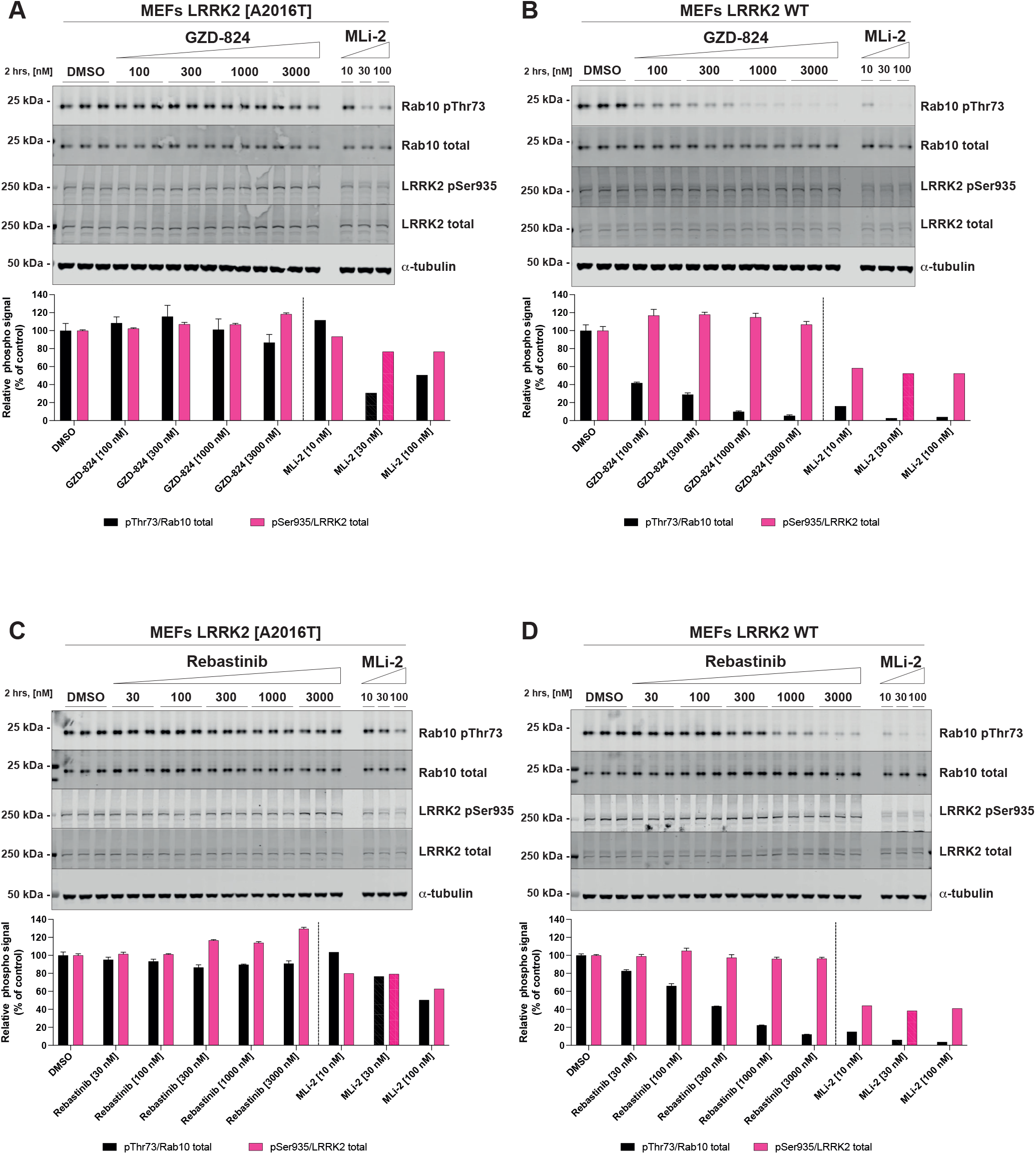
GZD-824 and Rebastinib do not induce Rab10 Thr73 dephosphorylation in inhibitor resistant LRRK2 [A2016T] cells. (A+C) Inhibitor resistant immortalized MEFs LRRK2 [A2016] were treated with or without the indicated concentrations of inhibitors for 2 h. Cells were lysed, and 20 µg of extract was subjected to quantitative immunoblot analysis with the indicated antibodies (all at 1 µg/ml). Each lane represents cell extract obtained from a different dish of cells (three replicates per condition). (B+D) As in (A+C) except that the cells were wild type. The membranes were developed using the Odyssey CLx Western Blot imaging system. Immunoblots were quantified using the Image Studio software. Data are presented relative to the phosphorylation ratio observed in cells treated with DMSO (no inhibitor), as mean ± SD. (A+B) SD are derived from the replicates show in the presented blots.

### GZD-824 does not inhibit 14-3-3-binding or Rab29 mediated LRRK2 recruitment to the Golgi

Phosphorylation of Ser910 and Ser935 regulates the ability of LRRK2 to interact with 14-3-3 adaptor proteins [11]. To study the impact that GZD-824 has on 14-3-3 binding, we treated HEK293 cells overexpressing wild type LRRK2 with 5 μM GZD-824 for 4 h, immunoprecipitated LRRK2 and immunoblotted for co-immunoprecipitated endogenous 14-3-3 using an antibody that picks up multiple human isoforms. We observed that GZD-824 had no significant impact on 14-3-3 binding under these conditions, whereas 1 μM GSK3357679A ablated 14-3-3 binding and Ser935 phosphorylation (Fig 7A).

**Figure 7.**
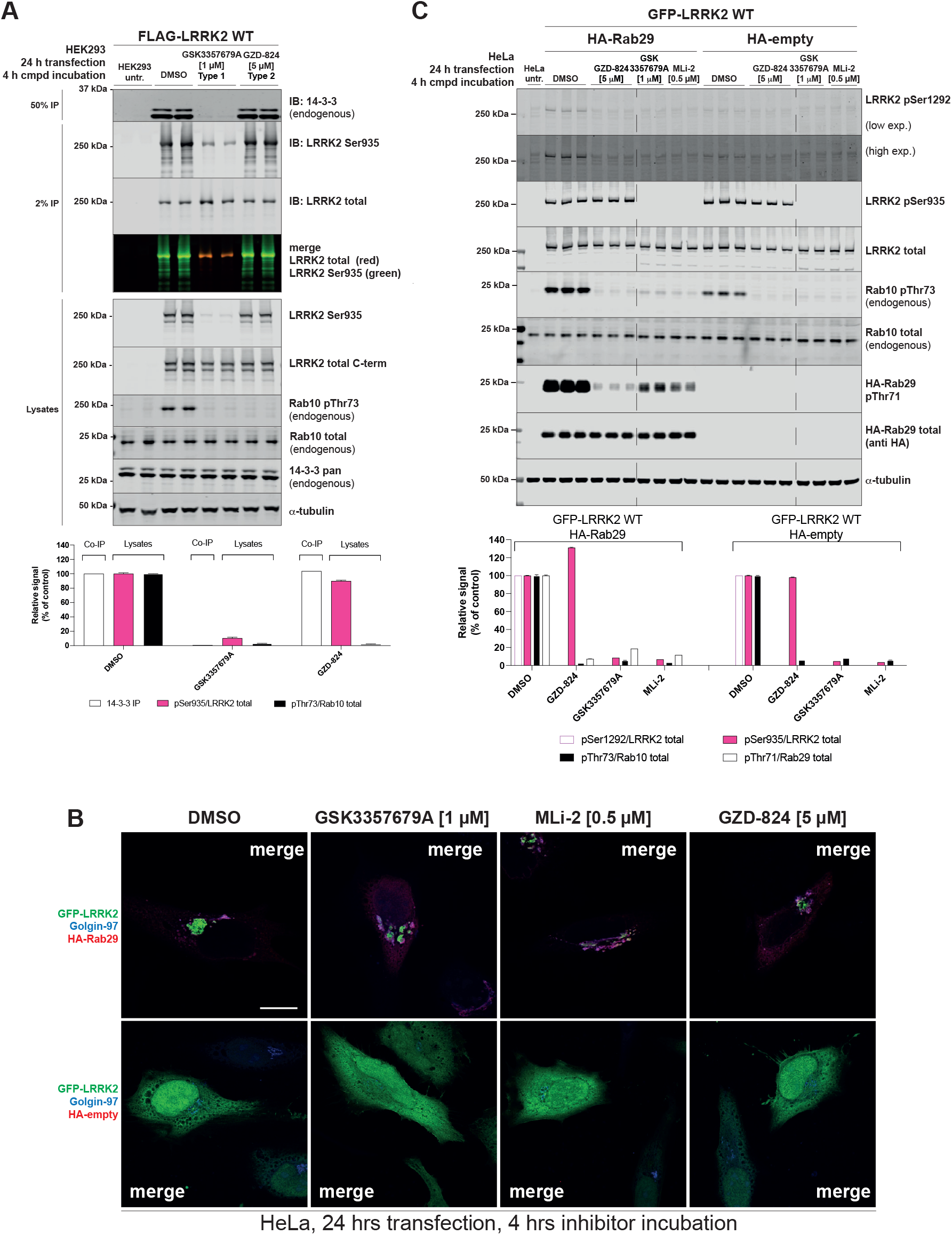
GZD-824 has no impact on 14-3-3 and Rab29 binding. (A) HEK293 cells were transiently transfected for 24 h with the indicated constructs. 24 h post-transfection cells were treated with the indicated concentrations of inhibitors for the indicated times. 15 µg of whole cell extracts was subjected to quantitative immunoblot analysis with the indicated antibodies (all at 1 µg/ml). Flag-LRRK2 is immunoprecipitated and subjected to immunoblotting with the indicated antibodies. Each lane represents cell extract obtained from a different dish of cells. The membranes were developed using the Odyssey CLx Western Blot imaging system. (B) HeLa cells were transiently transfected with wild type GFP-LRRK2 in the presence of absence of HA-Rab29. 24h post-transfection cells were treated ± the indicated concentration of inhibitor for 4 h. Cells were fixed in 3.7% (by vol) paraformaldehyde and stained with mouse anti-HA and anti-Golgin-97 (trans Golgi marker). Scale bar represents 10 μm. The Figures are representative of at least two independent experiments. (C) HeLa cells were transiently transfected for 24 h with the indicated constructs. 24 h post-transfection cells were treated with the indicated concentrations of inhibitors for the indicated times. 15 µg of whole cell extracts was subjected to quantitative immunoblot analysis with the indicated antibodies (all at 1 µg/ml). Each lane represents cell extract obtained from a different dish of cells. The membranes were developed using the Odyssey CLx Western Blot imaging system (A+C) Immunoblots were quantified using the Image Studio software. Data are presented relative to the phosphorylation ratio observed in cells treated with DMSO (no inhibitor), as mean ± SD. (A+C) SD are derived from the replicates shown in the presented blots.

Rab29 is localized at the Golgi and co-expression of LRRK2, and Rab29 in HeLa cells strikingly results in the bulk of LRRK2 being recruited to the Golgi apparatus (Fig 7B, SFig3-4) [56]. Treatment of HeLa cells co-expressing LRRK2 and Rab29 with 5 μM GZD-824 for 4 h did not significantly impact Rab29 mediated relocalization of LRRK2 to the Golgi apparatus (Fig 7B). In parallel experiments, treatment with 1 μM GSK3357679A for 4h also had no impact on recruitment of LRRK2 to the Golgi by Rab29 (Fig 7B). Control immunoblotting study for this experiment is shown in Fig 7C. Consistent with previous work [56], the LRRK2 mediated phosphorylation of endogenous Rab10 at Thr73 and autophosphorylation at Ser1292 is enhanced by overexpression of Rab29 (Fig 7C). Both Type I and Type II inhibitors suppress LRRK2 autophosphorylation at Ser1292, as well as phosphorylation of Thr73 of endogenous Rab10. Type II inhibitors do not impact LRRK2 Ser935 phosphorylation. We also show that LRRK2 mediated phosphorylation of Rab29 at Thr71, monitored using a recently released commercial phospho-specific antibody, is blocked by both Type I and Type II inhibitors (Fig 7C).

### GZD-824 stimulates basal mitophagy similarly to GSK3357679A

Previous work revealed that pathogenic LRRK2[G2019S] mutation significantly reduces basal mitophagy, but not general autophagy, in primary MEFs and various mouse tissues in a manner that is rescued by inhibiting LRRK2 with structurally distinct and selective Type I inhibitors, GSK3357679A and MLi-2 [25]. We therefore monitored the impact that 10-300 nM GZD-824 had on basal mitophagy in previously generated wild type, LRRK2[G2019S] knock-in and LRRK2 knock-out MEFs [7] stably expressing the *mito*-QC mitophagy reporter [25]. The *mito*-QC reporter is localized to mitochondria via an outer mitochondrial membrane targeting sequence derived from the protein FIS1 and bears a tandem GFP and mCherry tag. When a mitochondrion is delivered to lysosomes, the low lysosomal luminal pH quenches the GFP but not the mCherry signal. The degree of mitophagy is determined by the appearance of mCherry-only puncta, which represent mitochondria engulfed into autolysosomes [57-59]. Treatment of wild type or LRRK2[G2019S] cells for 24 h with 30-300 nM GZD-824 stimulated basal mitophagy ∼2-fold, to the same extent as GSK3357679A (Fig 8A, 8B). This effect was specific for inhibition of LRRK2, as neither GZD-824 or GSK3357679A impacted basal mitophagy in LRRK2 knock-out cells. Consistent with previous work [25], basal mitophagy was lower in LRRK2[G2019S] MEFs compared to wild type MEFs and treatment with Type I or Type II inhibitors increased basal mitophagy to levels similar to those observed in LRRK2 knock-out cells (Fig 8A, 8B).

**Figure 8.**
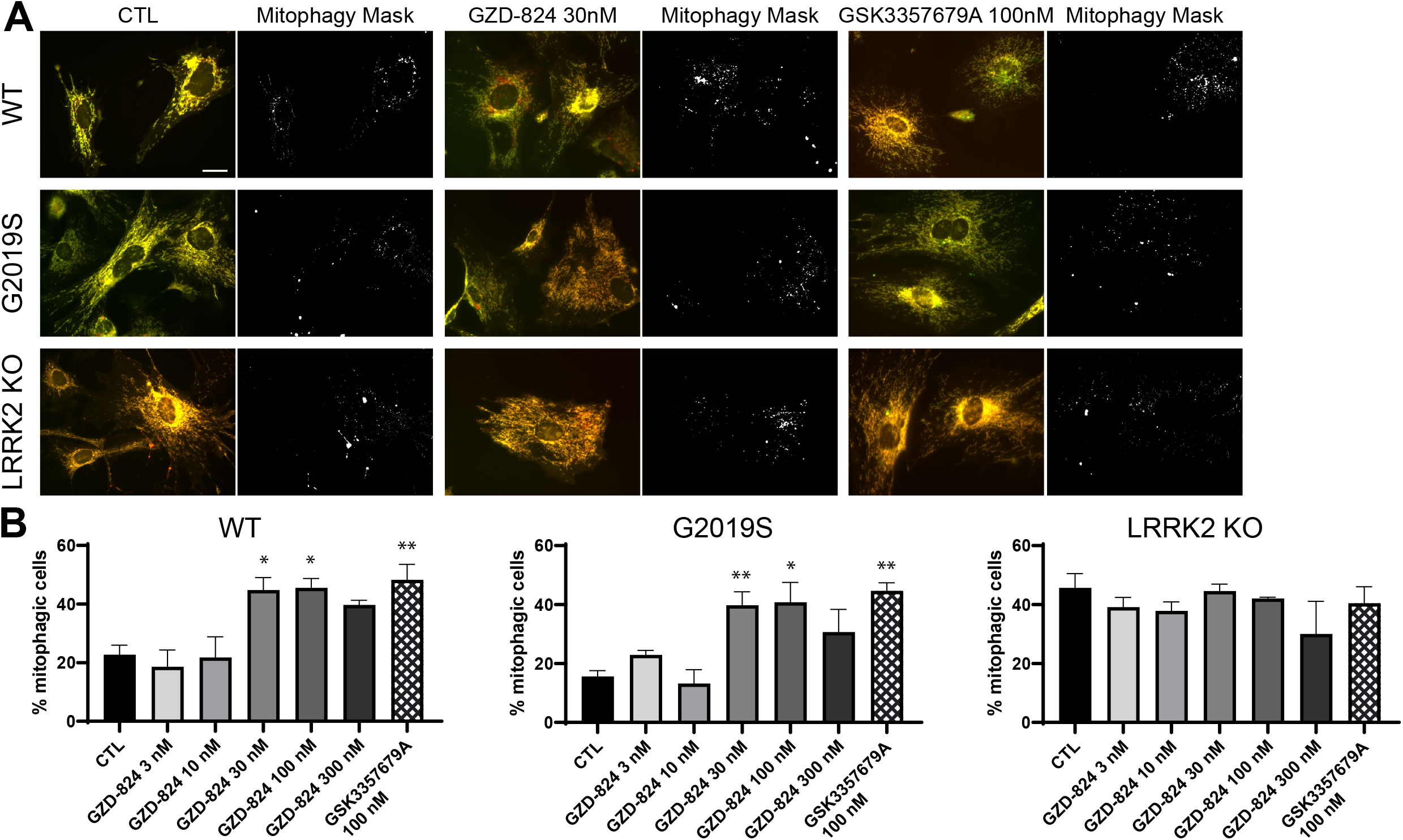
GZD-824 enhances mitophagy in primary MEFs. (A) Representative images and corresponding mitophagy mask generated with the mito-QC counter [55] of wild-type, homozygous LRRK2 G2019S, and LRRK2 KO primary mito-QC MEFs. Cells were treated with or without the indicated concentrations of inhibitors for 24 h. (B) Quantitation of mitophagy in cells treated with incremental doses of GZD-824. Data are represented as the mean ± SEM of 3-4 experiments and was analysed with a one-way ANOVA followed by a Dunnett’s multiple comparison test. *p<0.05, **p,0.01.

## Discussion

In this study we compared the in vitro and cellular properties of Type I and 3 distinct Type II LRRK2 inhibitors. All Type II inhibitors studied, suppressed LRRK2 protein kinase activity in cells as judged by their ability to inhibit LRRK2 mediated Rab protein phosphorylation and stimulate basal mitophagy similar to Type I compounds. The most striking difference between Type I and Type II compounds is that Type II inhibitors do not induce dephosphorylation of Ser935 or other LRRK2 biomarker phosphorylation sites. This suggests that Type II inhibitors preserve biomarker site phosphorylation on LRRK2 by maintaining the LRRK2 kinase domain in the open inactive conformation. The mechanism underlying this phenomenon is not known, but we hypothesize that in the Type II-open kinase domain conformation, the N-terminal biomarker sites on LRRK2 are exposed for phosphorylation by the upstream kinase(s) and/or rendered inaccessible to protein phosphatase(s). Vice versa, in the Type I-closed conformation, access of the N-terminal biomarker sites to upstream kinase(s) could be restricted/or N-terminal biomarker sites could be better exposed to protein phosphatase(s). In future analysis it would be important to compare how Type I and Type II inhibitors influence the conformation and accessibility of the biomarker sites to upstream kinases and phosphatases.

The finding that prolonged treatment of cells with GZD-824 does not impact on Ser935 phosphorylation under conditions where Rab10 phosphorylation remains suppressed, argues against the biomarker sites located in the N-terminal region of LRRK2 being controlled by autophosphorylation. Although distinct kinases have been proposed to phosphorylate these sites [60-62], further work is required to validate which kinase(s) target these residues in vivo. A protein phosphatase-1 (PP1) complex has been reported to dephosphorylate these sites in vivo [63]. Future work should also explore whether interaction of LRRK2 with microtubule is involved in controlling biomarker phosphorylation. Targeting of LRRK2 to microtubules in the closed kinase conformation, could bring LRRK2 in association with a microtubule associated phosphatase and/or restrict access to an upstream protein kinase.

Our conclusions are also consistent with the finding that the majority of pathogenic mutations that promote the active closed conformation of LRRK2 (R1441G/C, Y1699C, I2020T) suppress biomarker site phosphorylation [11]. Consistent with this, treatment of LRRK2[R1441C] knock-in MEFs with GZD-824 (Fig 4) or Rebastinib (Fig 5), would be predicted to induce the open conformation and thus rescue Ser935 phosphorylation and this is what we observe. Our results emphasize that the steady state phosphorylation of the biomarker sites on LRRK2 correlates with the open vs closed conformation of the LRRK2 kinase domain rather than the intrinsic kinase activity towards Rab substrates. The finding that treatment of cells with Type II inhibitors does not further increase biomarker site phosphorylation suggests that the bulk of the cellular LRRK2 is in the open inactive conformation.

Neither GZD-824 (Fig S1A), Rebastinib (Fig S1B) or Ponatinib (Fig S1C) are selective kinase inhibitors. Our finding that LRRK2[A2016T] mutation suppresses inhibition of LRRK2 by GZD-824 as well as Rebastinib (Fig 6), could be exploited in future studies to better distinguish whether physiological effects of these inhibitors are mediated by LRRK2, or other cellular kinases inhibited by these compounds. When testing physiological effects of the Type II compounds used in this study, we would also recommend verifying that the cellular effects are observed with diverse Type II LRRK2 inhibitors as off targets of these inhibitors differ to some extent (Fig S1). Ponatinib is clinically approved for treating chronic myeloid leukemia and Philadelphia chromosome–positive acute lymphoblastic leukemia [64]. Rebastinib is currently being explored in Phase-1 clinical trials as an TIE2 kinase inhibitor for advanced or metastatic solid tumors (https://clinicaltrials.gov/ct2/show/NCT03717415). GZD-824, Rebastinib and Ponatinib inhibit wild type LRRK2 more potently than pathogenic LRRK2 in vitro (Table 1) and in cells (Fig 4, 5), a feature that will likely be similar for other Type II inhibitors. As pathogenic mutations favor the closed conformation of LRRK2 kinase domain, it would be expected that a higher concentration of a Type II inhibitor would be required to inhibit pathogenic LRRK2. This difference in potency between wild type and pathogenic LRRK2 for Type II inhibitors would need to be taken into consideration when evaluating the optimal therapeutic doses of a potential future Type II inhibitor compound for treating LRRK2 driven Parkinson’s disease.

To our knowledge all of the well-studied LRRK2 inhibitors induce dephosphorylation of the N-terminal biomarker sites on LRRK2 suggesting that these inhibitors will be Type I compounds. Previous work in LRRK2-deficient rodents as well as nonhuman primates treated with LRRK2 inhibitors highlighted preclinical liability concerns in lung and kidney, resulting from loss of LRRK2 kinase activity that might be associated with altered autophagy and lysosome biology [65-67]. In future work it would be important to explore the physiological impact that selective Type II LRRK2 inhibitors have on animal models in comparison to Type I compounds. It is possible that Type II compounds that do not induce microtubule association or lead to dephosphorylation of biomarker sites may have a different physiological impact. We would argue that the poor selectivity of GZD-824, Rebastinib and Ponatinib precludes their use as advanced tool compounds to probe and compare the detailed physiological impacts of Type I and Type II LRRK2 inhibitors. In future work we would advocate that it would therefore be important to develop more selective Type II LRRK2 tool inhibitors to enable their properties to be better studied.

## Acknowledgements

We thank Paul Davies for helpful discussion, Esther Sammler for preparing human neutrophils and the excellent technical support of the MRC-Protein Phosphorylation and Ubiquitylation Unit (PPU) DNA Sequencing Service (coordinated by Gary Hunter), the MRC-PPU tissue culture team (coordinated by Edwin Allen), MRC PPU Reagents & Services antibody and protein purification teams (coordinated by Hilary McLauchlan and James Hastie) and kinase profiling (coordinated by Jennifer Moran) and Natalia Shpiro for generating MLi-2 and preparation of Figure 1.

## Data availability

All of the primary data that is presented in this study can be requested in electronic form by contacting Dario Alessi and will also be deposited on the Zenodo data repository (link available after journal acceptance). All plasmids and antibodies (and associated datasheets) generated at the MRC Protein Phosphorylation and Ubiquitylation Unit at the University of Dundee can be requested through our reagent’s website https://mrcppureagents.dundee.ac.uk/. Requests for provision of GSK3357679A should be directed to Alastair Reith.

## Author Contributions

A.T designed, executed all experiments in this study presented in Figures 2, 3, 4, 5, 6, 7 Supplementary Figure 2A, 3, 4 analyzed and interpreted data and preparation of the manuscript. F.S. designed and executed and interpreted experiments relating to mitophagy analysis in Fig 8. I.G.G. oversaw studies relating to autophagy analyses. A.D.R was involved with experimental design, analysis, interpretation of data and preparation of the manuscript. D.R.A. helped with experimental design, analysis and interpretation of data and preparation of the manuscript.

## Figure and Table legends

**Supplementary Figure 1.**
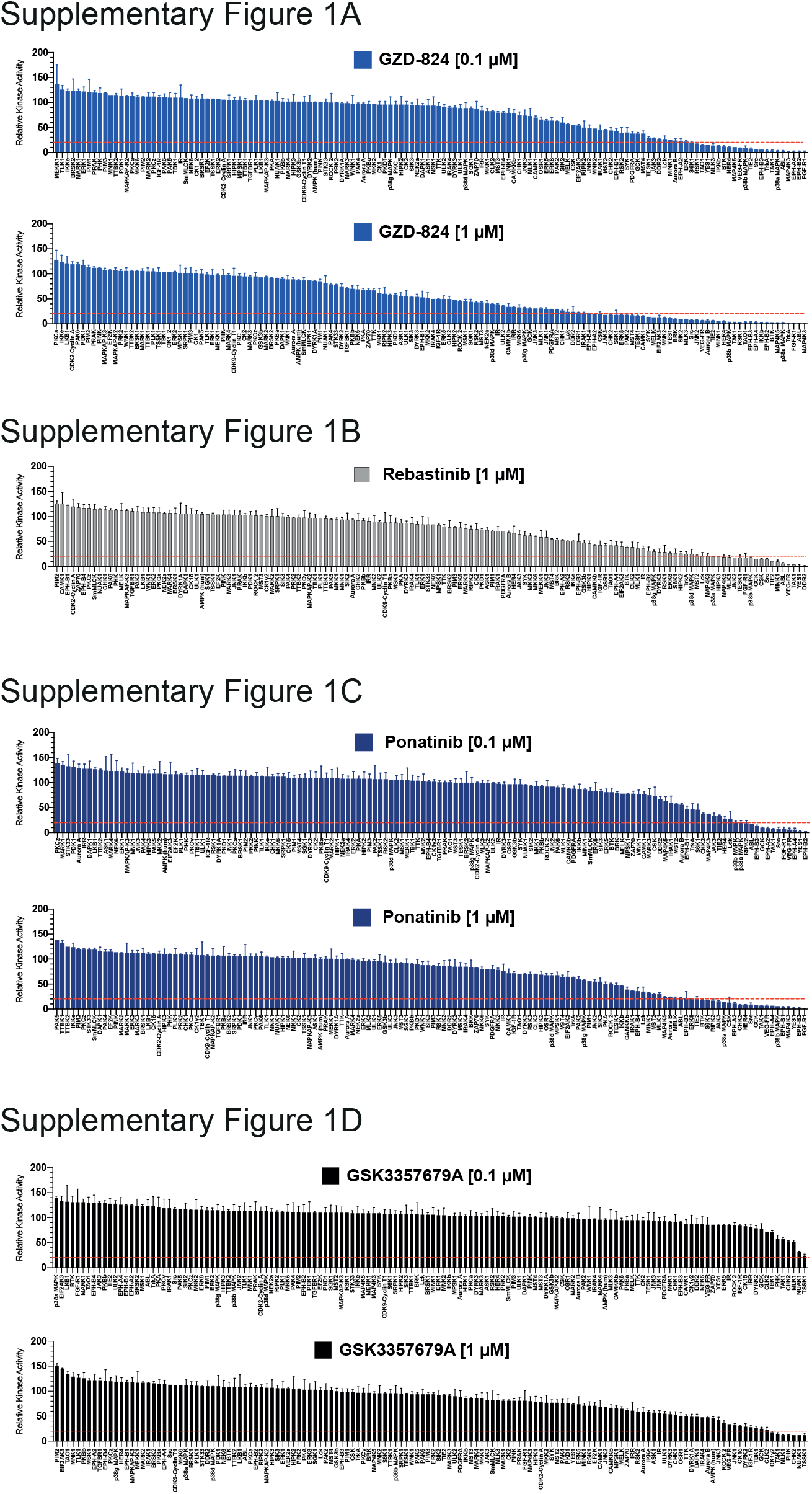
Kinase profiling of GZD-824, Ponatinib and GSK3357679A. Kinase profiling of GZD-824 (A), Rebastinib (B), Ponatinib (C) and GSK3357679A (D) were undertaken at a concentration of 0.1 and 1 μM against the Dundee panel of 140 protein kinases at the International Centre for Protein Kinase Profiling (http://www.kinase-screen.mrc.ac.uk). Results for each kinase are presented as the mean kinase activity± S.D. for an assay undertaken in duplicate relative to a control kinase assay in which the inhibitor was omitted. Abbreviations and assay conditions used for each kinase are available (http://www.kinase-screen.mrc.ac.uk/services/premier-screen).

**Supplementary Figure 2.**
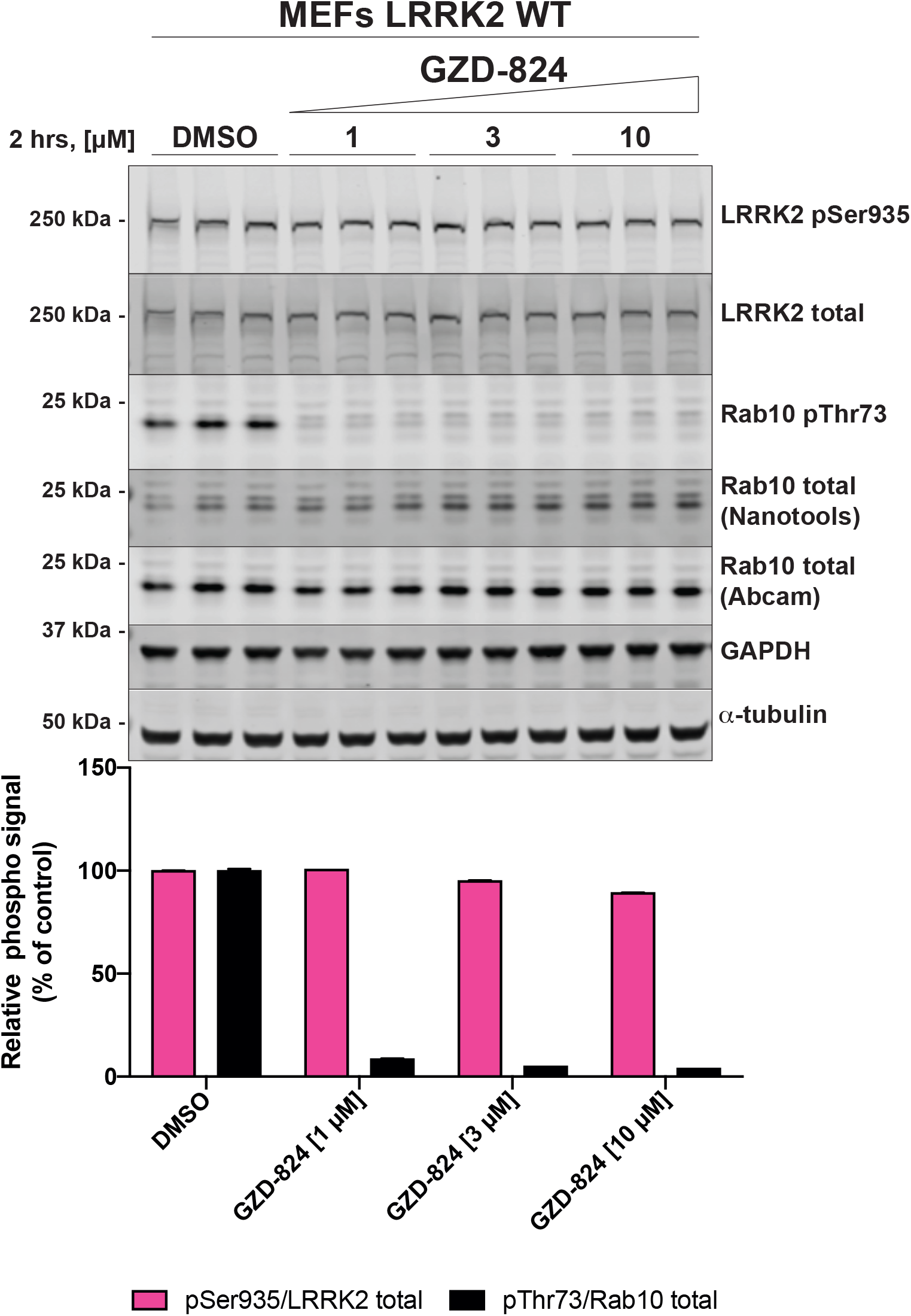
(A) Wild-type MEFs were treated with or without the indicated concentrations of inhibitors for 2 h. Cells were lysed, and 20 µg of extract was subjected to quantitative immunoblot analysis with the indicated antibodies (all at 1 µg/ml). Each lane represents cell extract obtained from a different dish of cells (three replicates per condition). The membranes were developed using the Odyssey CLx Western Blot imaging system. Immunoblots were quantified using the Image Studio software. Data are presented relative to the phosphorylation ratio observed in cells treated with DMSO (no inhibitor), as mean ± SD. SD are derived from the replicates shown in the presented blot.

**Supplementary Figure 3.**
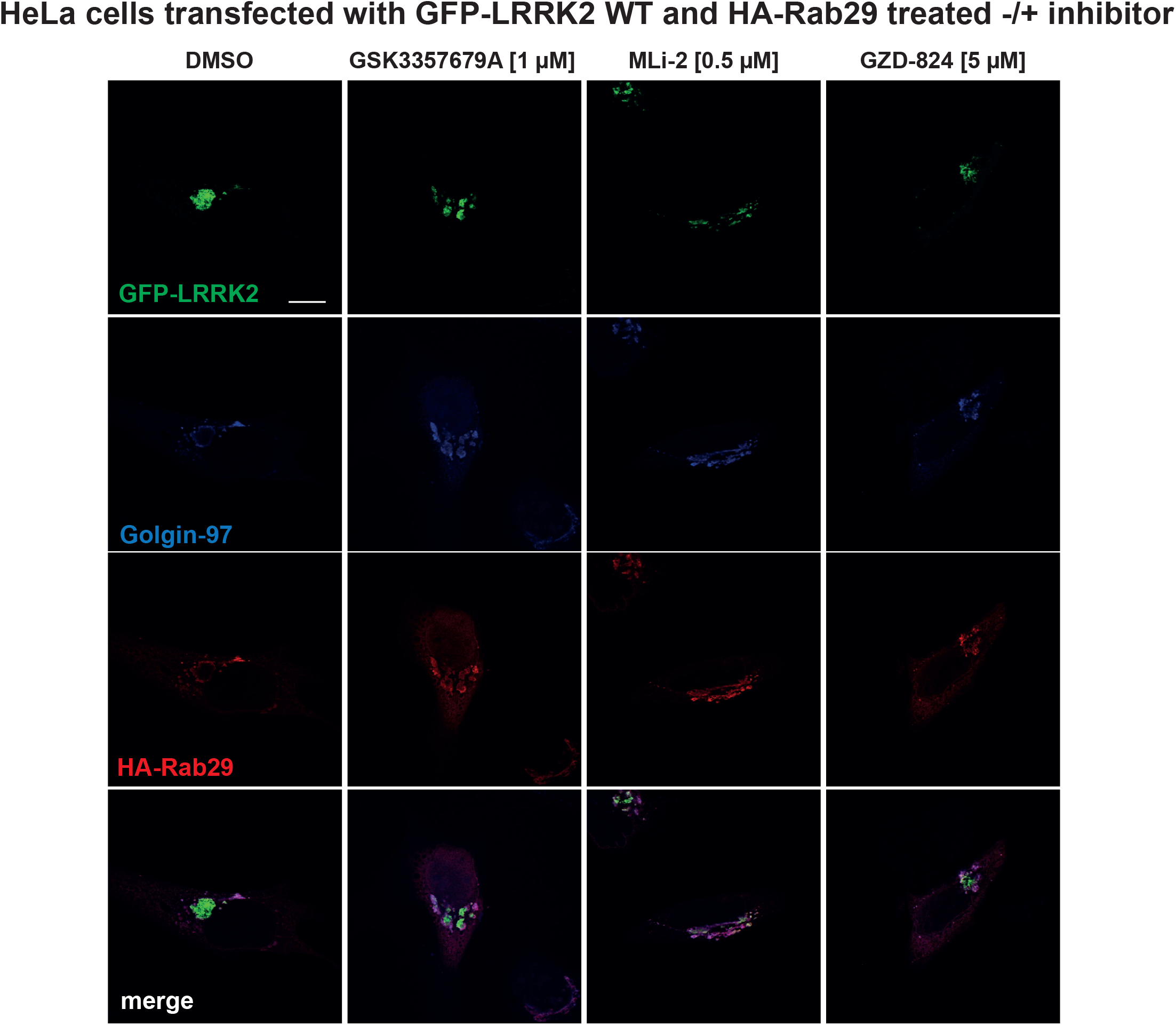
Control immunofluorescence study that accompanies Figure 4B. HeLa cells were transiently transfected with wild type GFP-LRRK2 in the presence of HA-Rab29. 24h post-transfection cells were treated ± the indicated concentration of inhibitor for 4 h. Cells were fixed in 3.7% (by vol) paraformaldehyde and stained with mouse anti-HA and anti-Golgin-97 (trans Golgi marker). Scale bar represents 10 μm. The Figures are representative of at least two independent experiments.

**Supplementary Figure 4.**
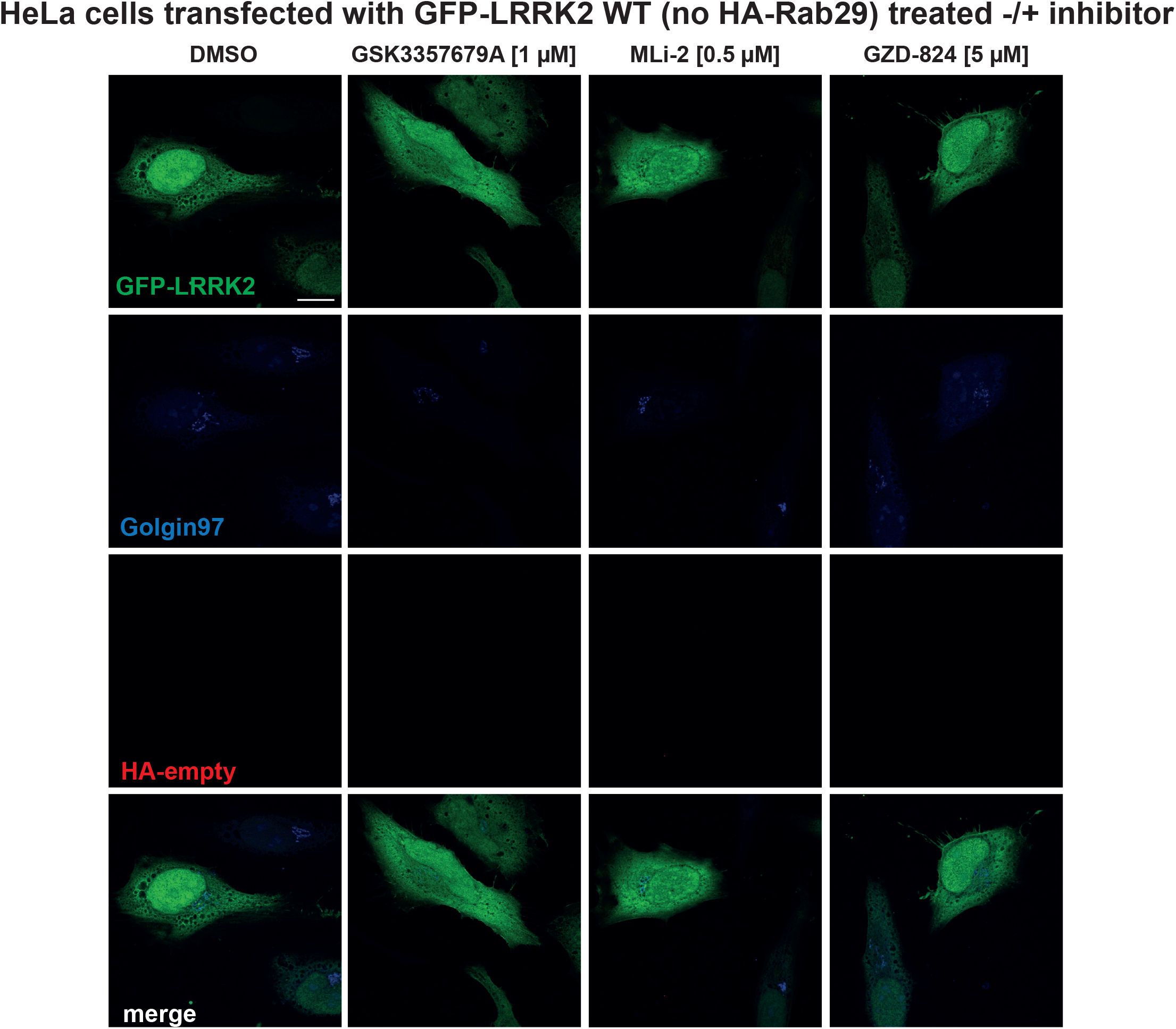
Control immunofluorescence study that accompanies Figure 4B. HeLa cells were transiently transfected with wild type GFP-LRRK2 in absence of HA-Rab29. 24h post-transfection cells were treated ± the indicated concentration of inhibitor for 4 h. Cells were fixed in 3.7% (by vol) paraformaldehyde and stained with mouse anti-HA and anti-Golgin-97 (trans Golgi marker). Scale bar represents 10 μm. The Figures are representative of at least two independent experiments.

